# Large-scale voltage imaging in the brain using targeted illumination

**DOI:** 10.1101/2021.04.05.438451

**Authors:** Sheng Xiao, Eric Lowet, Howard J. Gritton, Pierre Fabris, Yangyang Wang, Jack Sherman, Rebecca Mount, Hua-an Tseng, Heng-Ye Man, Jerome Mertz, Xue Han

## Abstract

Recent improvements in genetically encoded voltage indicators enabled optical imaging of action potentials and subthreshold membrane voltage dynamics from single neurons in the mammalian brain. To perform high speed voltage imaging, widefield microscopy remains an essential tool for recording activity from many neurons simultaneously over a large anatomical area. However, the lack of optical sectioning makes widefield microscopy more prone to background signal contamination, and thus far voltage imaging using fully genetically encoded voltage indicators remains limited to simultaneous sampling of a few cells over a restricted field-of-view. We here demonstrate a strategy for large scale voltage imaging using the fully genetically encoded voltage indicator SomArchon and targeted illumination. We implemented a simple, low-cost digital micromirror device based targeted illumination strategy to restrict illumination to the cells of interest, and systematically quantified the improvement of this microscopy design theoretically and experimentally with SomArchon expressing neurons in single layer cell cultures and in the brains of awake mice. We found that targeted illumination, in comparison to widefield illumination, increased SomArchon signal contrast and reduced background cross-contamination in the brain. Such improvement permitted the reduction of illumination intensity, and thus reduced fluorescence photobleaching and prolonged imaging duration. When coupled with a high-speed, large area sCMOS camera, we routinely imaged tens of spiking neurons simultaneously over minutes in the brain. Thus, the widefield microscopy design with an integrated targeted illumination system described here offers a simple solution for voltage imaging analysis of large neuron populations in behaving animals.

## Introduction

Recent advances in genetically encoded voltage indicators (GEVIs) have enabled neuroscientists to directly measure membrane voltage from individual neurons in the mammalian brain^1–6^. In particular, a few recent GEVIs, including SomArchon, QuasAr3, Voltron, ASAP3 and Ace2N have achieved sufficient sensitivity to capture individual action potentials from single neurons recorded from behaving mice. Of these high performance GEVIs, several are fully genetically encoded, whereas others are hybrid sensors that require exogenous chemicals^7–13^. One class of fully genetically encoded indicators detects voltage dependent fluorescence of fluorophores fused to voltage sensitive peptide domains derived from voltage gated ion channels, voltage sensitive phosphatases or rhodopsins^7–12^. For these GEVI designs, changes in cell membrane voltage induce confirmational transitions of the voltage sensitive domains, which subsequently alter the intensity or the efficiency of Forster resonance energy transfer of the tethered fluorophores. A recent example is ASAP3 that measures voltage dependent fluorescence of a circular permutated GFP fused to the voltage sensing domain of G. gallus voltage-sensing phosphatase^4^. Another class of fully genetically encoded indicators are single compartment and directly detect the intrinsic voltage dependent fluorescence of engineered rhodopsins, such as QuasAr3, Archon and SomArchon^3,5,14^. To improve fluorescence signals, bright chemical fluorophores have also been explored in the designs of GVEIs, yielding a class of high-performance hybrid GEVIs that requires both exogeneous chemical dyes and the corresponding voltage sensing protein counterparts^2,13,15^.

With rapid and continued improvements of GEVIs, voltage imaging offers great promise for direct analysis of neuronal voltage dynamics in the brain. To capture fast membrane voltage fluctuations, especially action potentials that occur on the millisecond and sub-millisecond time scale, fluorescence voltage imaging needs to be performed at a near kilohertz sampling speed. Point scanning techniques, such as multiphoton microscopy, have minimum signal cross-contamination and out-of-focus background due to confined excitation volumes^16^, but are generally limited to video-rate acquisition speed as they rely on mechanical scanners. Fast random access scanning using acousto-optic deflectors has been demonstrated with kilohertz sampling rates^4^, although these devices require a complicated setup, are sensitive to motion artifacts, and more importantly, can only record very few pre-selected cells at once. More recently, kilohertz frame rate two-photon imaging over a field-of-view (FOV) of 50 × 250 µm^2^ has been demonstrated by means of passive pulse splitting from a specialized low-repetition rate laser^17^. However, its stringent alignment requirements, high cost, and concerns regarding long-term system stability remain a major obstacle for its widespread use for neuroscience studies.

Alternatively, widefield microscopy, especially when equipped with the newly developed high-speed large-area sCMOS cameras, remains a cost-effective and easily implementable solution for wide FOV, kilohertz frame-rate imaging. This ability to image a large FOV at high spatiotemporal resolution is particularly critical to resolving morphological details of individual neurons, and to correct for tissue movement associated with physiological processes (i.e., heart rate, breathing) that are unavoidable when imaging the brains of awake behaving animals. However, a major limitation of widefield microscopy is the inability to reject out-of-focus and scattered light^18^, making it prone to signal contamination and background shot noise caused by non-specific excitations^19^. To address this problem molecularly, recently developed GEVIs have utilized soma targeting peptides to restrict the expression of GEVIs to the soma or the proximal dendrites. For example, SomArchon, QuasAr3, Voltron, and ASAP3-kv all include the axon initial segment targeting motif of the potassium channel Kv2.1^20^, which are critical for their success in the brains of behaving animals ^2–6^. Restricting the expression of GEVIs to a sparse subset of neurons can also help reduce background signal contamination *in vivo*, and this strategy was recently used to achieve simultaneous imaging of tens of neurons using the hybrid sensor Voltron^2^.

In parallel with the molecular targeting of GEVIs, targeted illumination has also been developed to enhance image contrast and signal-to-noise ratio (SNR) ^5,6,21^. By using a digital micromirror device (DMD) or a spatial light modulator to pattern the illumination light, targeted illumination confines fluorescent excitation to pre-selected areas of interest based on reference images previously obtained from the same sample. It has been implemented in extended-depth-of-field microscopy for video rate calcium imaging^21^, and was recently demonstrated to significantly improve voltage imaging performance with a widefield microscope^5,6^. However, for voltage imaging applications, this approach thus far has been limited to simultaneous sampling of a few cell, and requires sophisticated microscopy setups.

We integrated a simple, low cost DMD-based targeted illumination module into a standard widefield microscope, and directly compared SomArchon imaging performance of the same neurons under targeted versus widefield illumination, in both cultured neuron preparations, and from the brains of awake behaving mice. We found that illumination targeting reduced nonspecific background fluorescence and fluorescence signal cross-contamination, leading to increased SomArchon spike signal-to-background ratio. The improvement of SomArchon fluorescence contrast allowed us to decrease the total excitation power over the imaging FOV with reduced fluorescence decay. As a result, we were able to perform routine SomArchon voltage imaging from tens of neurons over a large anatomical area of 360 × 180 µm^2^ while maintaining SomArchon fluorescence contrast, and over a prolonged recording duration of several continuous minutes. These results demonstrate that targeted illumination with a DMD represents a simple, low cost, and practical strategy for large scale voltage imaging of tens of neurons over an extended period of time in awake behaving animals.

## Results

### Modeling and testing the effects of targeted illumination on optical crosstalk in widefield optical imaging

Motivated by the unique advantage of widefield microscopy in performing optical voltage imaging with high spatiotemporal resolution over large FOVs, we considered a targeted illumination approach to further enhance signal quality by reducing out-of-focus background signals. We first developed a theoretical model to estimate how targeted illumination might minimize signal crosstalk due to out-of-focus excitation or tissue scattering from nearby neurons that are not actively being imaged. We considered contributions from both out-of-focus fluorescence and tissue scattering, by modeling light propagation through scattering media using the radiative transfer equation in the forward scattering limit^22^ (see Methods and Fig. S1). In our model, we characterized crosstalk values from non-targeted neurons at distance *D*_*x*_ laterally and *D*_*z*_ axially from a region of interest (ROI) under widefield versus targeted illumination conditions (Fig. 1a, b, and Fig. S3). We found that in simulated fluorescence images, targeting illumination to a specific neuron substantially reduced the overall background in the imaging plane, and therefore reduced the strength of crosstalk from neighboring neurons (red circle: Fig. 1d, e). Additionally, the level of crosstalk contamination from a non-overlapping axially displaced neuron is strongly affected by the distance between the out-of-focus neurons relative to the imaging plane (Fig. 1c). These findings confirm that, for widefield microscopy, targeting illumination to a neuron of interest can improve signal quality by reducing the overall fluorescence background, and limiting signal contamination from neighboring out-of-focus neurons. These computational results highlight that targeted illumination is a viable approach for low-background, high-contrast imaging of voltage signals in the brain using widefield microscopy.

**Figure 1:**
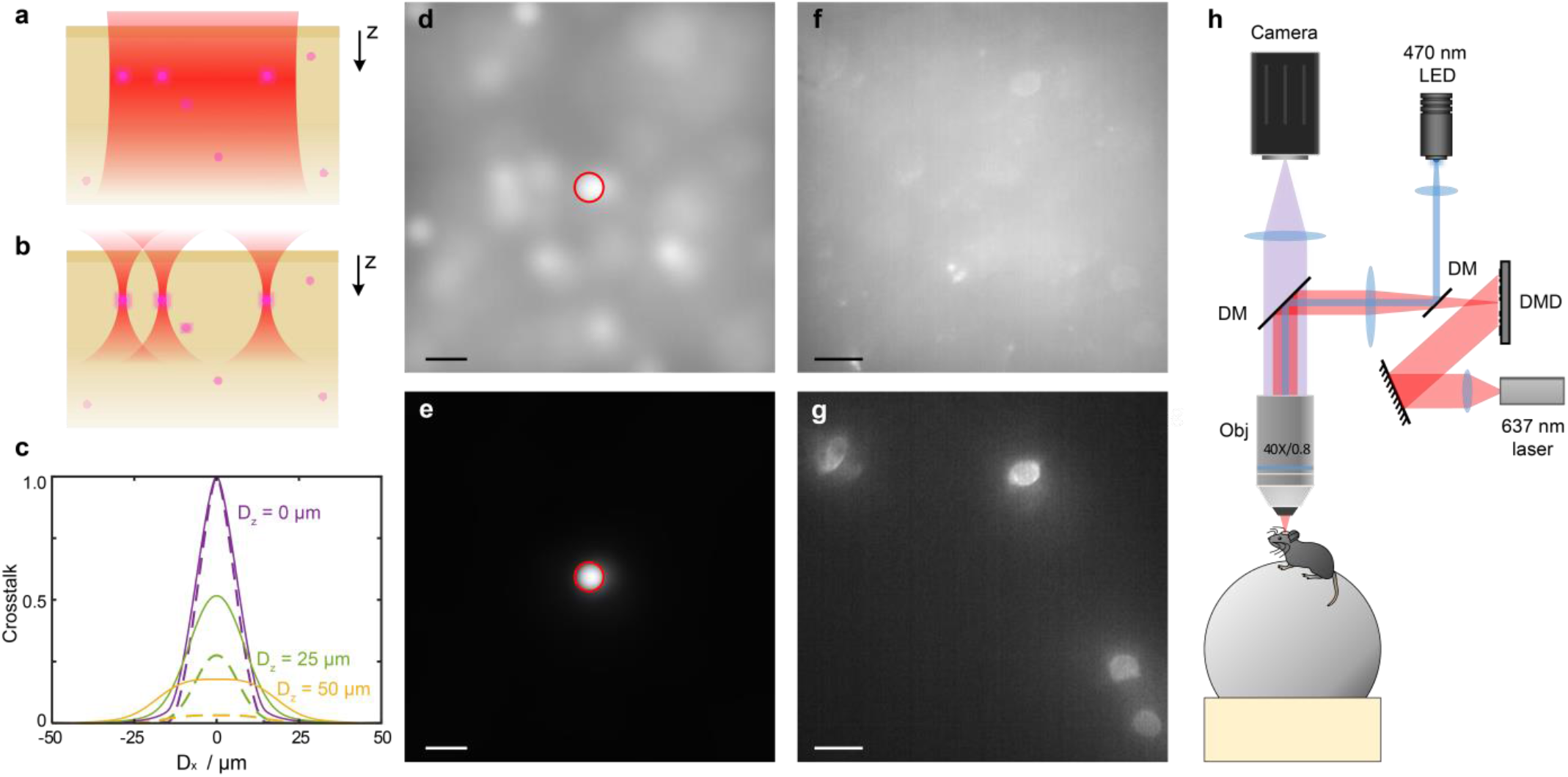
Theoretic models of targeted illumination on fluorescence crosstalk in widefield optical imaging and a microscopy design for experimental testing in awake mice. (**a, b**) Illustration of the theoretical consideration of fluorescence imaging of individual neurons using widefield illumination (**a**) and targeted illumination (**b**). Purple dots illustrate the location of individual SomArchon expressing neurons. Red area illustrates the illumination beam for SomArchon excitation. (**c**) Characterization of fluorescence crosstalk values from a non-targeted neuron at a lateral distance D_x_ and axial distance D_z_ away from the neuron of interest under widefield and targeted illumination conditions. Solid lines, widefield illumination; dashed line, targeted illumination. Purple, D_z_ = 0 µm; green, D_z_ = 25 µm; yellow, D_z_ = 50 µm. (**d**,**e**) Simulated images of fluorescence from a single neuron under widefield illumination (**d**) and targeted (**e**) illumination. Illumination target is indicated by the red circle. (**f, g**) An example widefield versus targeted illumination voltage imaging experiment in an awake head fixed mouse positioned on a spherical treadmill shown in (**h**). SomArchon fluorescence of visual cortex neurons imaged with widefield illumination (**f**), and targeted illumination (**g**) of 4 individual neurons under the same laser power density. (**h**) Experimental setup. DM, dichromatic mirror; Obj, objective lens. Scale bars are 20 µm.

To experimentally evaluate the improvement of targeted illumination, we integrated a DMD into a custom-built widefield microscope configured for dual-color GFP and SomArchon imaging (Fig. 1h). We performed voltage imaging of SomArchon expressing neurons, in both cell cultures and in the visual cortex and the hippocampus of awake head fixed mice. Since SomArchon protein is fused to the GFP reporter, static GFP fluorescence images were first taken to identify SomArchon expressing neuronal soma. The GFP fluorescence images were then used to generate references for targeting illumination to the identified SomArchon positive neurons. Consistent with what was observed in our computational models, we found that restricting illumination to the soma reduced the overall background fluorescence and accordingly enhanced the contrast of SomArchon fluorescence in individual cells (Fig. 1f, g).

### Targeted illumination increases spike spike-to-baseline (SBR) ratio and reduces SomArchon photobleaching in cultured neurons

We first examined whether targeted illumination improves SomArchon voltage imaging quality in cultured neurons transduced with AAV9-syn-SomArchon. Cultured neurons on flat glass coverslips have little out-of-focus fluorescence originating from the out of plane z-axis, and therefore should only exhibit small amount of signal contamination (Fig. S2). To directly compare the effects of targeted versus widefield illumination, we alternated 20-second long imaging trials between the two illumination conditions for the same FOV (n = 226 neurons recorded from 16 FOVs).

Cultured neurons exhibit spontaneous subthreshold membrane voltage fluctuations that occasionally produce action potentials. Since SomArchon can detect subthreshold voltage dynamics^3^, the actual photon shot noise is mixed with real biological subthreshold voltage fluctuations. Therefore, it is difficult to quantify the actual noise level and therefore accurately evaluate SomArchon signal qualities by calculating the absolute SNR. We thus calculated the spike signal-to-baseline ratio (SBR), defined as the amplitude of the spike divided by the variance of the experimentally measured baseline fluctuations (Vm), as an estimated performance metric of SomArchon in recording individual spikes. The estimation of Vm variation however is affected by the presence of suprathreshold spike events. We therefore developed a spike-insensitive SBR estimation method that does not require prior identification of spikes (see Methods). To test the properties of the SBR algorithm, we simulated membrane voltage using the Izhikevich-type neuron model that exhibit both action potentials and biological subthreshold voltage (Fig. S4a). Additionally, to model experimentally measured noise signals, we added different levels of Gaussian white noise (Fig. S4b, d, e). We calculated the theoretical SBR as spike amplitude divided by the experimentally measurable Vm that contains both the biological subthreshold voltage and the white noise. We further calculated the theoretical SNR as the spike amplitude divided by the variance of the white noise only. We found that our SBR estimation substantially underestimated the theoretical SNR, but it better reflected the theoretic SBR that considers biological Vm variation (Fig. S4c). Thus, though the spike SBR measure is an underestimation of SomArchon molecular performance, it provides an intuitive and spike-insensitive measure of the optical voltage signal quality, especially for in vivo recordings where conditions can vary substantially.

With targeted illumination, we detected a spike SBR of 5.7 ± 2.0 (mean ± standard deviation, from 226 neurons in 16 FOVs), significantly greater than that observed from the same neurons under the widefield illumination condition (4.9 ± 1.4, Fig. 2h). Since the spike identification algorithm relies on a custom spike SBR threshold, we investigated whether the increase in spike SBRs depends on the threshold used to identify spikes. We found that across several chosen SBR threshold values, targeted illumination consistently resulted in greater spike SBRs than widefield illumination (Fig. S5, Table S3). Furthermore, targeted illumination resulted in more detected spikes than that detected from the same neurons measured in the widefield condition (Fig. 2i).

**Figure 2.**
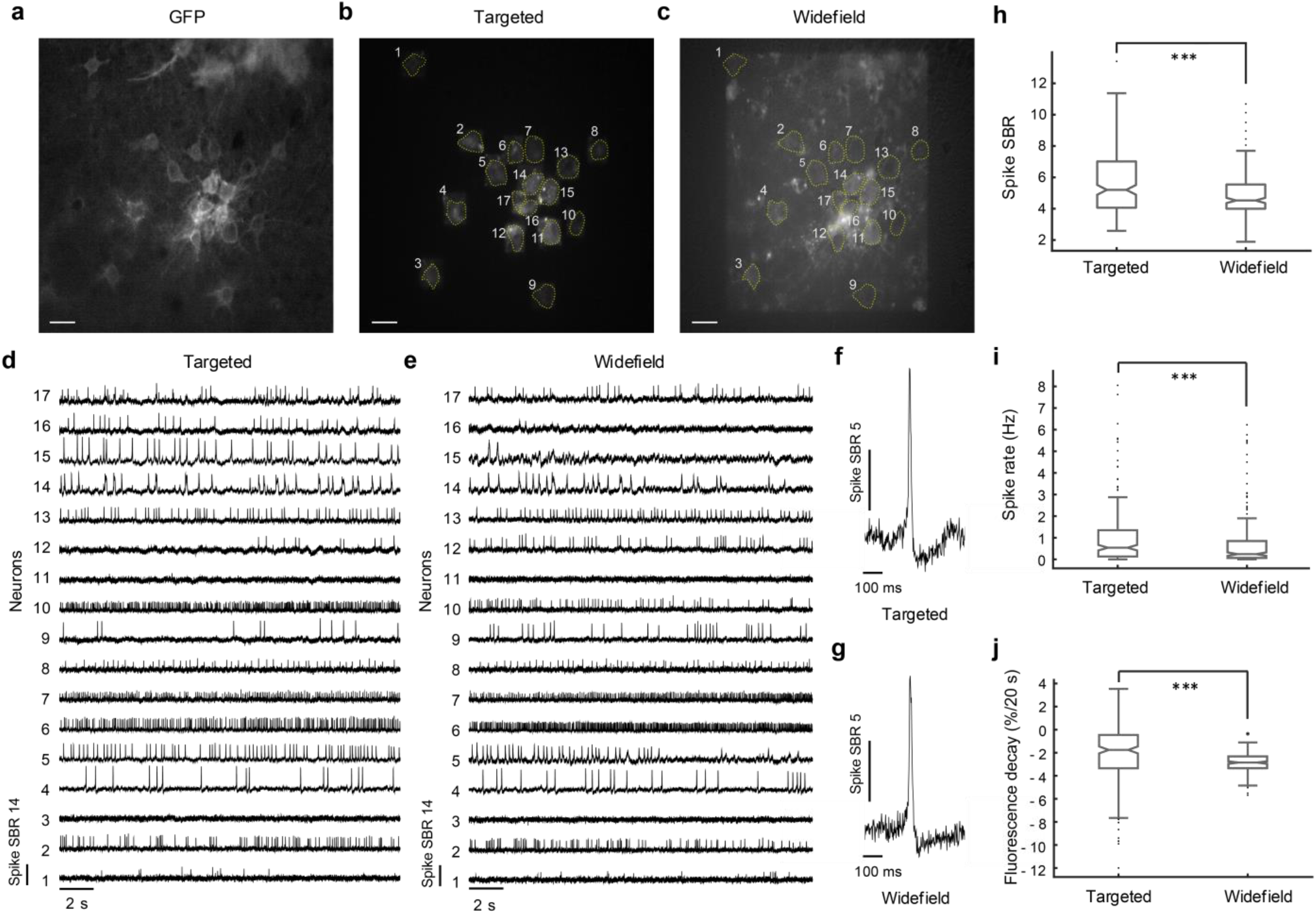
Targeted illumination increases spike SBR and reduces fluorescence decay in cultured neurons. (**a-c**) An example FOV showing cultured neurons expressing SomArchon fused to a static GFP fluorophore; scale bars are 20 μm. (**a**) GFP fluorescence image under widefield illumination. (**b**) SomArchon fluorescence under targeted illumination. (**c**) SomArchon fluorescence under widefield illumination. (**d, e**) Example SomArchon fluorescence traces from 17 simultaneously recorded neurons in the FOV illustrated in A, using targeted illumination (**d**), and widefield illumination (**e**). (**f, g**) Example individual spikes recorded from the same neuron with targeted illumination (**f**) and with widefield illumination (**g**). (**h**) Spike SBR (***, p = 6.73e^-14^, paired t-test comparing targeted illumination versus widefield illumination conditions, df = 204, n = 226 neurons from 16 FOVs). (**i**) Spike rate identified with a spike SBR threshold of 4.5 (***, p = 4.23e^-5^, paired t-test, df = 225). (**j**) Reduction of SomArchon fluorescence over 20 seconds period (***, p = 9.06e^-7^, paired t-test, df = 225). The illumination power density was ∼2 W/mm^2^ for both the target illumination and the widefield illumination conditions, for all recordings from cultured neurons. For all boxplots, the box indicates the median (middle line), 25th (Q1, bottom line), 75th (Q3, top line) percentiles, and the whiskers are Q1-1.5*(Q3-Q1), and Q3+1.5*(Q3-Q1). Outliers that exceed these values are shown as dots.

We next examined fluorescence decay, calculated as the percent reduction of fluorescence intensity over time, an important parameter that limits the duration of fluorescence imaging in general. We found that with targeted illumination, SomArchon showed a slight fluorescence decay of 2.15 ± 2.66% (mean ± standard deviation, n = 226 neurons) over a 20 second recording period, significantly smaller than that observed under widefield illumination (2.99 ± 1.04%, Fig. 2j). Together, these results demonstrate that targeted illumination significantly improves SomArchon performance in terms of spike SBR and fluorescence decay, even in cultured neurons where out-of-focus background is minimal.

### Targeted illumination increases the cross-correlation of spikes, but minimally impacts cross-correlation of subthreshold membrane voltage (Vm) in neuron cultures

With a high speed, large field of view sCMOS camera, we were able to image 2 - 28 neurons simultaneously (14.13 ± 7.59, mean ± standard deviation, from 16 FOVs) at 500 Hz over a FOV of 360 × 180 μm^2^. To estimate how targeted illumination can reduce signal crosstalk, we calculated the cross-correlation between neuron pairs, for both Vm and spikes. We found that both Vm-Vm and spike-spike correlation decreased slightly with increasing anatomical distance between simultaneously recorded neuron pairs (slopes for linear regression between Vm-Vm correlation and distance are -5.3e^-4^ and -3.7e^-4^ for targeted illumination and widefield illumination respectively, and between spike-spike correlation and distance are -1.7e^-4^ and -7.2e^-5^ respectively, Fig. 3a,b). The regression slopes of Vm-Vm correlation and spike-spike correlation over anatomical distance under the targeted illumination condition are both greater than that observed during the widefield illumination condition (Vm-Vm correlation, p = 5.3592e^-6^, z score = -4.5502, permutation test; spike-spike correlation, p = 0.0039, z score = -2.8865).

**Figure 3.**
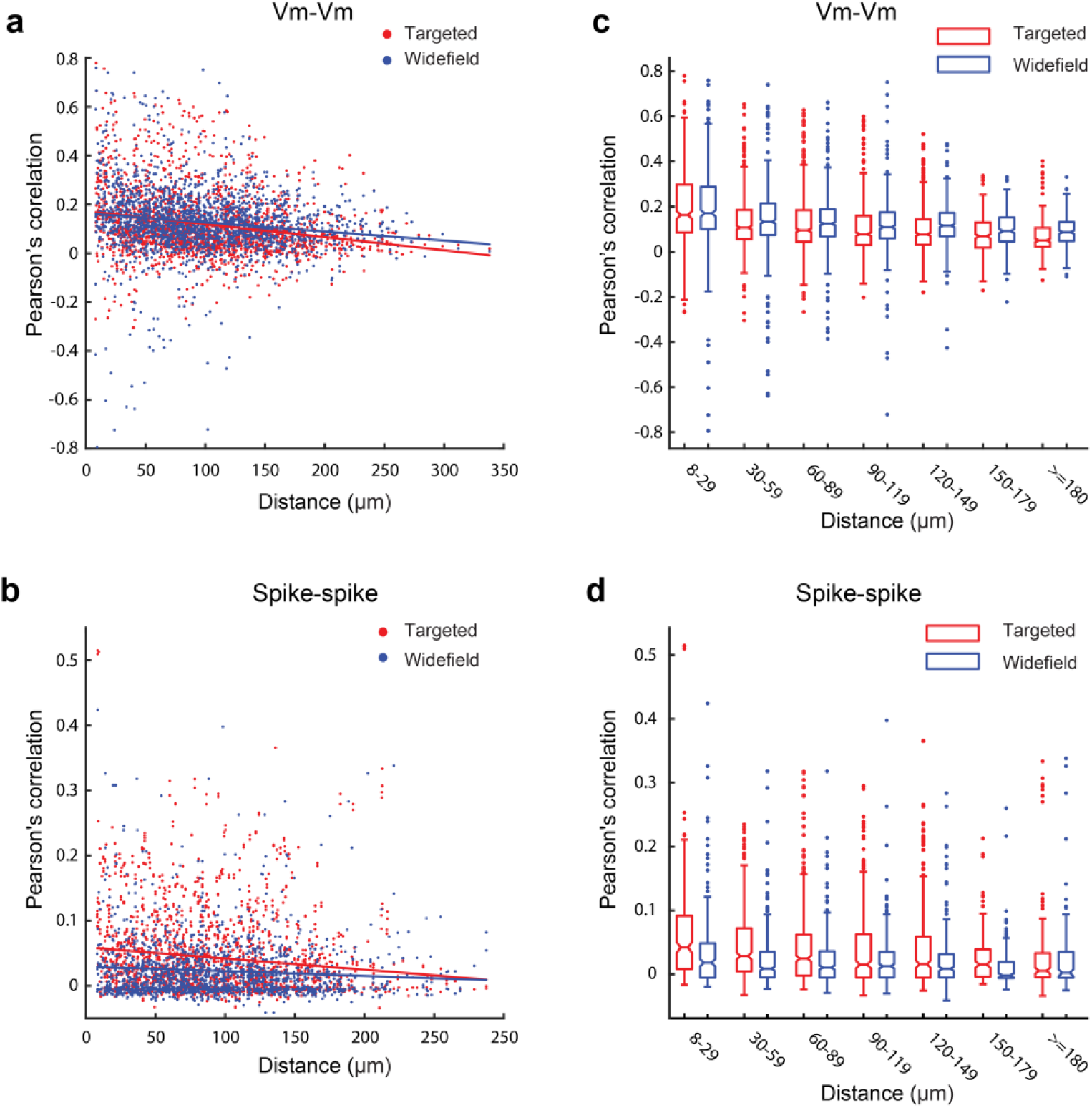
Targeted illumination effects on Vm-Vm correlation and spike-spike correlation in cultured neurons. (**a**,**b**) Pearson’s correlation values between pairs of simultaneously recorded neurons decreased over anatomical distance for Vm-Vm correlation (**a**) and spike-spike correlation (**b**). Red dots indicate correlation values from pairs of neurons recorded under the targeted illumination condition. Blue dots indicate correlation values from pairs of neurons recorded under the widefield illumination condition. (**c, d**) Vm-Vm correlation (**c**) and spike-spike correlation (**d**) at different distances with 30 µm increment. Red boxplots are correlation values obtained with the targeted illumination condition, and blue boxplots are with the widefield illumination condition. For all boxplots, the box indicates the median (middle line), 25th (Q1, bottom line), 75th (Q3, top line) percentiles, and the whiskers are Q1-1.5*(Q3-Q1), and Q3+1.5*(Q3-Q1). Outliers that exceed these values are shown as dots. Refer to Tables S1 and S2 for statistical tests.

To further evaluate changes in Vm-Vm and spike-spike correlation across different anatomical distances between the two illumination conditions, we binned the correlation values of neuron pairs every 30 µm. Consistent with the improvement of spike SBR observed under the targeted illumination condition, spike-spike correlation was slightly greater under targeted illumination than widefield illumination condition across neuron pairs within 180 µm, although no difference was observed for neuron pairs over 180 µm (Fig. 3d). When we examined Vm-Vm correlation, we found no difference between targeted illumination and widefield illumination conditions for neurons pairs within 120 µm, though a slightly smaller correlation value was obtained under targeted illumination for neurons over 120 µm away (Fig. 3c). The similar Vm-Vm correlations under widefield and targeted illumination is consistent with our numerical models when the sample is only a monolayer of cells absent of significant contributions from out-of-focus and scattered fluorescence (Fig. S2).

### Targeted illumination improves SomArchon spike SBR, reduces fluorescence crosstalk, and enables long-duration recording in the visual cortex of awake mice

To quantify the effect of targeted illumination in the brains of awake animals, we examined SomArchon expressing neurons in the superficial layers of visual cortex. Mice were head-fixed and able to freely locomote on a spherical treadmill. For each FOV, we alternated 10-second long voltage imaging sessions between targeted illumination and widefield illumination conditions. We found that targeted illumination significantly reduced the decay of SomArchon fluorescence (11.02 ± 3.08% over 10 seconds), approximate half of that observed with widefield illumination (20.38 ± 3.05%, p = 4.74^-14^, paired t-test, df = 20, Fig. 4k). Targeted illumination also resulted in a significant increase in SomArchon spike SBR, achieving 4.6 ± 0.7, significantly higher than the 4.1 ± 0.44 obtained with widefield illumination (p = 0.023, paired t-test, df = 18 neurons, Fig. 4l), and similar to that observed in cultured neurons (Fig. 2h). This significant increase in spike SBR for the targeted illumination condition accordingly led to a greater frequency of spikes identified when spike SBR threshold is used for spike identification (Fig. 4m).

**Figure 4:**
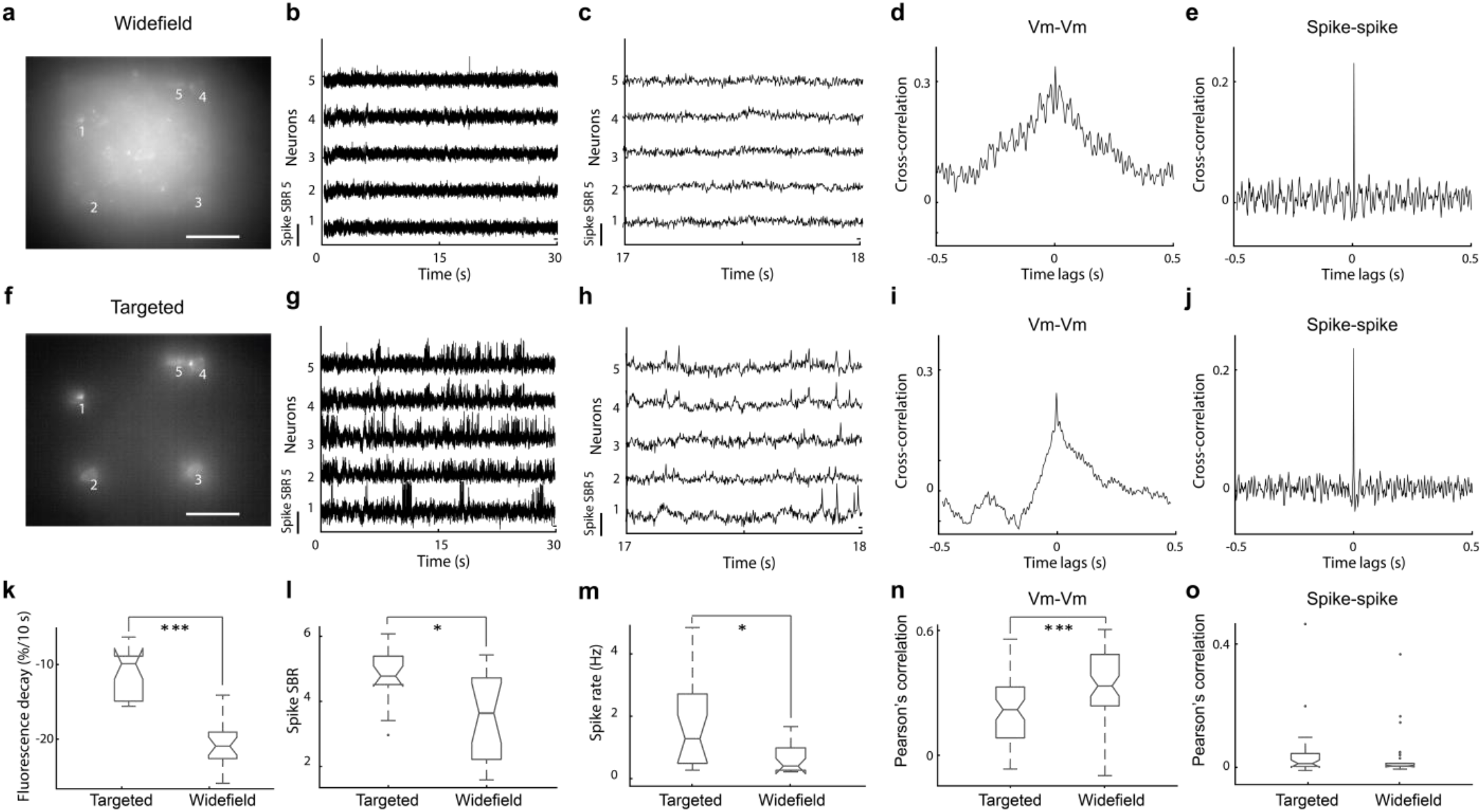
Targeted illumination improves SomArchon voltage imaging performance. (**a-e**) SomArchon fluorescence voltage imaging under widefield illumination condition. (**a**) An example FOV showing SomArchon fluorescence intensity averaged over the recording session. Scale bar: 50 µm. (**b**,**c**) Example SomArchon fluorescence traces (**b**), and zoomed in view (**c**), from simultaneously recorded 5 neurons indicated in (**a**) over a 10 second long recording period. (**d, e**) Example Vm-Vm (**d**) and spike-spike correlation. (**e**) Cross-correlogram of a single neuron pair, neurons labeled 2 and 3 in (**a**). (**f-j**) SomArchon fluorescence voltage imaging under targeted illumination condition. (**g-j**) Same plots and calculation as in (**b-e**), but for targeted illumination condition. (**k**) Fluorescence decay over 10 seconds for widefield vs targeted illumination conditions (***, p = 4.74e^-14^, paired t-test, df = 20). (**l**) Spike SBR for widefield vs targeted illumination conditions (*, p = 0.023, paired t-test, df = 18). (**m**) Detected spike rates (*, p = 0.016, paired t-test, df = 18). (**n**) Vm-Vm correlations between simultaneously recorded neuron pairs with targeted illumination versus widefield illumination (***, p = 0.00027, paired t-test, df = 30). (**o**) Spike-spike correlation between simultaneously recorded neuron pairs with targeted illumination versus widefield illumination (p = 0.58, paired t-test, df = 30). The illumination power density was ∼3 W/mm^2^ for both the target illumination and the widefield illumination conditions, for this and all other visual cortex recordings. For all boxplots, the box indicates the median (middle line), 25th (Q1, bottom line), 75th (Q3, top line) percentiles, and the whiskers are Q1-1.5*(Q3-Q1), and Q3+1.5*(Q3-Q1). Outliers that exceed these values are shown as dots.

To examine how targeted illumination impacts correlation measurements between simultaneously recorded neuron pairs, we computed spike-spike and Vm-Vm correlations as detailed above in cultured neuron experiments. Unlike in cultured neurons, here due to tissue scattering and fluorescence from out-of-focus neurons, we observed that targeted illumination significantly reduced Vm-Vm correlation values (Fig. 4n). However, spike-spike correlation values remained largely consistent under both conditions (Fig. 4o). Since spikes are only produced when Vm depolarization reaches sodium channel activation threshold for action potential generation, joint synaptic inputs that produce correlative low amplitude Vm changes between neuron pairs that are subthreshold in theory will not be captured by spike-spike correlation measures. The fact that Vm-Vm correlation is reduced by targeted illumination highlights that Vm signals contain a higher proportion of background signal crosstalk than spiking signals.

With targeted illumination, the drastically reduced background fluorescence and consequently enhanced spike SBR allowed us to reduce illumination intensity during voltage imaging. This reduction of overall ballistic illumination intensity, and the minimization of ROI exposure to backscattered light, can help reduce SomArchon photobleaching and thus allowed for recording over an extended duration under targeted illumination. Figure 5 represents an example of a continuous recording of 5 minutes, revealing that excellent spike SBRs can be maintained throughout the recording duration. Of the two simultaneously recorded neurons, the spike SBR for neuron 1 was 4.13 ± 1.1 (mean ± standard deviation, n = 1366 spikes), and for neuron 2 was 4.81 ± 1.29 (mean ± standard deviation, n = 261 spikes). However, we did notice a reduction in spike SBR over time (p = 9e^-14^, Kruskal-wallis, df = 1626 spikes, from 2 neurons combined), which reflects the effect of fluorescence photobleaching.

**Figure 5:**
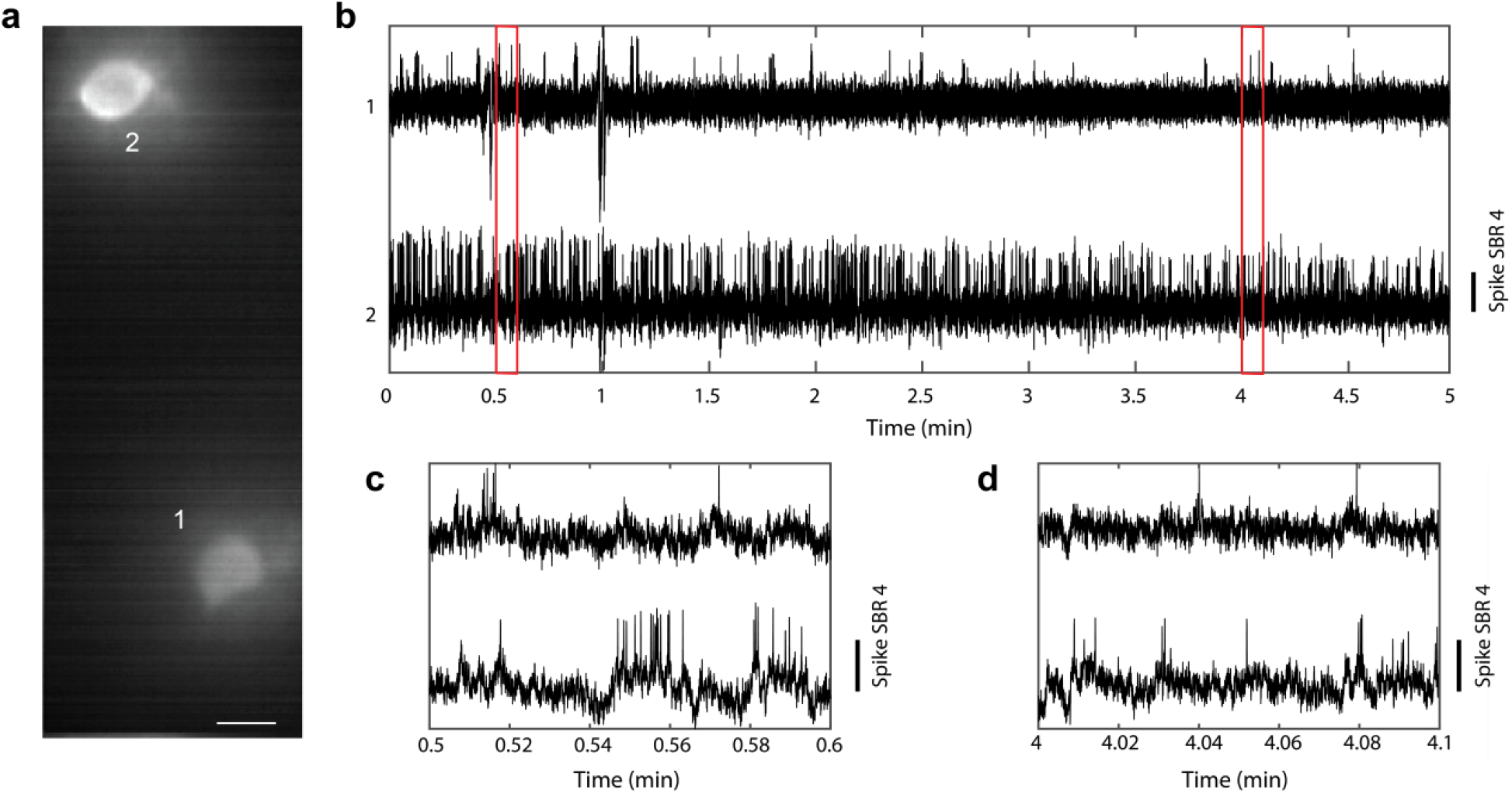
An example 5-minute long continuous recording session from two visual cortex neurons. (**a**) SomArchon fluorescence image from two neurons visualized with targeted illumination. Scale bar, 10 µm. (**b**) SomArchon fluorescence traces throughout the entire 5-minute long recording session. (**c**,**d**) Zoom-in view of SomArchon fluorescence towards the beginning (**c**) and the end of the recording session (**d**).

### Targeted illumination allows for large scale recordings in the hippocampal CA1 region in awake mice

Having established the significant advantage of targeted illumination, we deployed targeted illumination to image multiple neurons in the dorsal hippocampus CA1 region. Combining targeted illumination with a high-speed, large sensor sCMOS camera, we sampled a FOV of 360 × 180 µm^2^, often containing tens of neurons. We performed 6 recordings of 17 or more CA1 neurons (37 ± 22 neurons per session, mean ± standard deviation), while mice were awake and head-fixed navigating on a spherical treadmill. Across these recording sessions, we recorded a total of 222 spiking neurons, with a spike SBR of 4.16 ± 0.5 (mean ± standard deviation, n = 222 spiking neurons, Fig. 6, Fig. S6). In one recording, we were able to record 76 neurons simultaneously, and detected spikes in 58 of those neurons, over a 90-second long recording period (Fig. 6). The mean spike SBR of these neurons was 3.94 ± 0.4 (mean ± standard deviation, n = 58 neurons).

**Figure 6:**
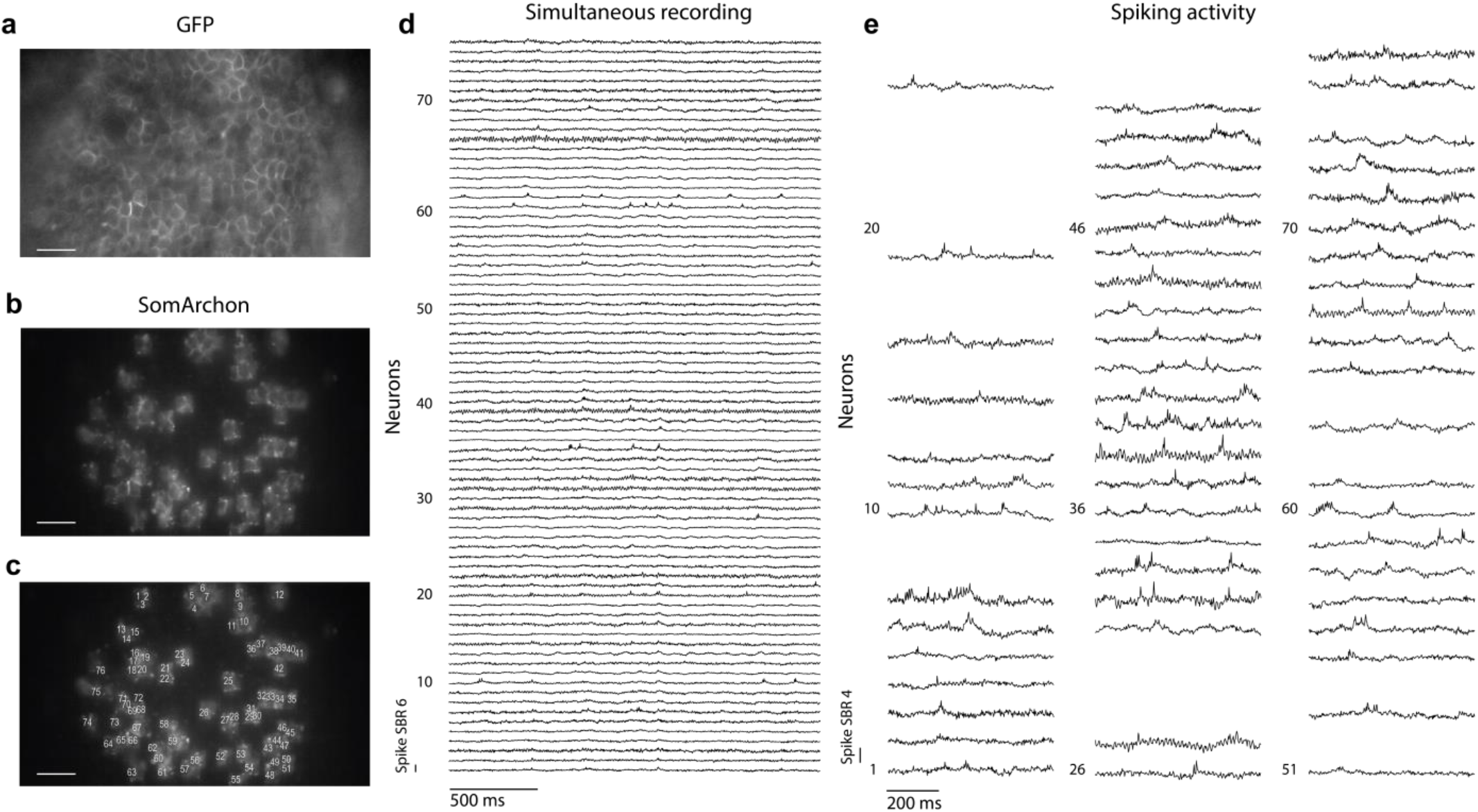
An example 90-second long continuous recording of 76 CA1 neurons simultaneously using targeted illumination in a behaving mouse. (**a-c**) SomArchon expressing CA1 neurons in the FOV, visualized via GFP fluorescence (**a**), SomArchon fluorescence visualized with targeted illumination (**b**), and with each neuron labelled (**c**). Scale bar, 50 µm. (**d**) Example traces of simultaneously recorded 76 CA1 neurons. 2.5 seconds recordings are shown here. (**e**) Representative example spikes in the 58 neurons where spikes were detected. The illumination power density was 4 - 5W/mm^2^ for this and all other CA1 recordings.

Further quantification of Vm-Vm and spike-spike correlation over anatomical distance between simultaneously recorded CA1 neuron pairs revealed that both Vm-Vm and spike-spike correlations substantially decreased with anatomical distance (Fig. 7, Kruskal-wallis, p = 5.16e^-7^, df = 5109 for Vm-Vm correlations; p = 3.18e^-6^, df = 4771 for spike-spike correlations). These results are consistent with that observed in cultured neurons and reflect the general understanding that nearby neurons tend to receive more temporally aligned synaptic inputs relative to neurons further apart^23^.

**Figure 7:**
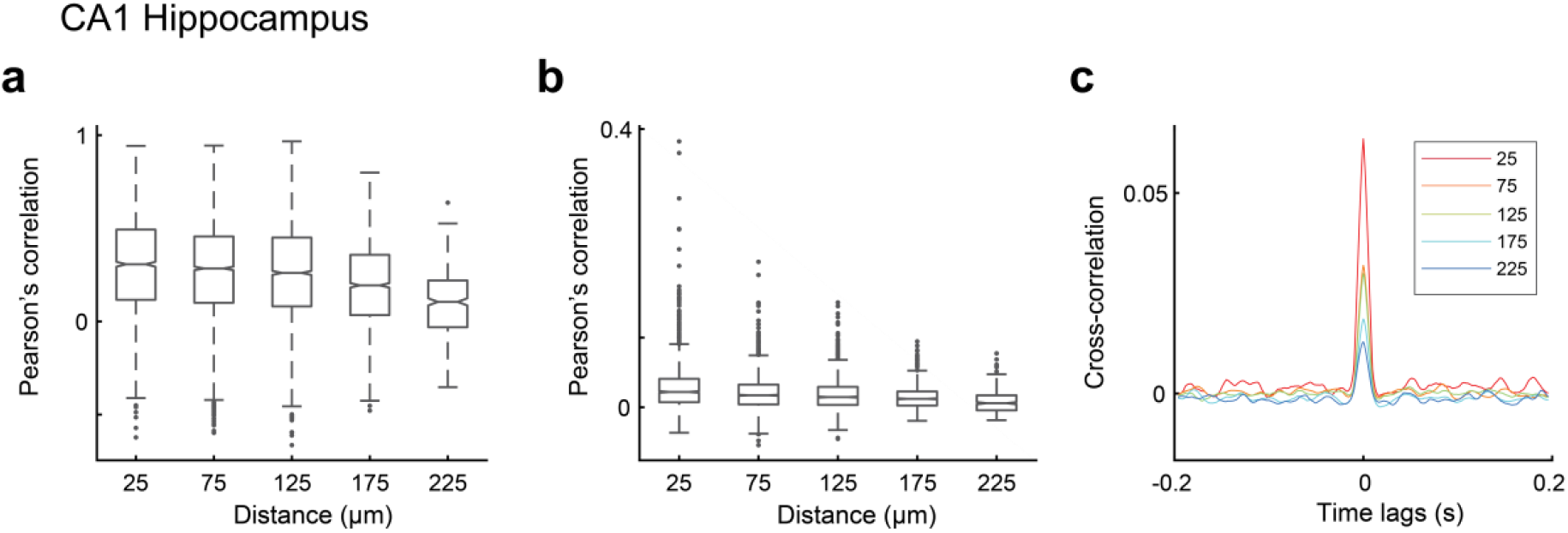
Pearson’s correlation values of spike-spike and Vm-Vm correlation over anatomical distance. (**a**) Vm-Vm correlation of CA1 neuron pairs over distance. Correlation values are grouped by distance (0 - 50 µm, 51 - 100 µm, 101- 150 µm, 151 - 200 µm, 201 - 250 µm). (**b**) Spike-spike correlation of CA1 neurons over distance. (**c**) Average spike-spike cross-correlogram across all recorded neuron pairs. For all boxplots, the box indicates the median (middle line), 25th (Q1, bottom line), 75th (Q3, top line) percentiles, and the whiskers are Q1-1.5*(Q3-Q1), and Q3+1.5*(Q3-Q1). Outliers that exceed these values are shown as dots.

## Discussion

Widefield microscopy remains an important imaging technique for high-speed voltage imaging, especially in task performing animals where a large FOV and a high spatial resolution are desired to resolve the activity from many individual neurons simultaneously. To improve widefield fluorescence microscopy for high speed, large scale, and long duration voltage imaging, we integrated a standard DMD-based targeted illumination system into a widefield microscope equipped with a high speed, large field of view sCMOS camera. This improved widefield microscopy design provides a simple, low-cost solution for large scale voltage imaging. We estimated the impact of background fluorescence on voltage imaging theoretically, and then experimentally quantified the improvement of targeted illumination for SomArchon voltage imaging in 2D neuron cultures and in the brains of behaving mice. We found that by restricting illumination to neuronal cell bodies, we were able to significantly increase SomArchon signal quality in terms of spike SBR and reduce out-of-focus background fluorescence. These improvements were more substantial for imaging neuron in the brain than in single layered neuron cultures, and were consistently observed across the two brain regions tested that have varying labeling density, including the superficial layers of visual cortex with sparsely labeled neurons and the hippocampus with densely labeled neurons. With such improvements in SomArchon signal quality, together with a high-speed large sensor size sCMOS camera, we were able to record optical voltage signals from over 70 neurons simultaneously over a wide FOV of 360 × 180 µm^2^ at 500 Hz.

One advantage of using targeted illumination is the reduced power density of excitation light, from both direct ballistic excitation photons and backscattered photons from tissue scattering. In this study, the ballistic excitation power density used for *in vivo* recordings was measured at 3 – 5 W/mm^2^, which equals to 0.7 – 1.1 mW per neuron (assuming a 15 x 15 µm^2^ square excitation region). However, for *in vivo* imaging, the actual excitation power will be further affected by tissue scattering. Photons targeting a cell can be scattered away from the ROI, whereas photons targeting non-ROI regions could eventually reach an ROI due to forward and backward scattering. Therefore, although the same excitation power density was applied in both targeted illumination and widefield illumination conditions, neurons under widefield illumination conditions were actually exposed to higher excitation power, increasing photobleaching therefore causing greater observed fluorescence decay. In addition, under targeted illumination, the reduced background and consequently improved spike SBR also allowed us to reduce the ballistic excitation power. As a result, we were able to perform continuous recordings over several minutes in duration, with only moderate reductions in spike SBR. While the performance of fluorescence based activity indicators are always limited by photobleaching, deploying trial-based study designs without excitation illumination during inter-trial-intervals should allow SomArchon to measure membrane voltage over many trials, and potentially over a greater cumulative period of time than demonstrated here using continuous illumination.

Other than large FOV and long-term imaging, our DMD-based targeted illumination widefield microscope also presents several additional advantages that would help to provide greater access to general voltage imaging applications. With the use of SomArchon, only a single-color excitation source is required to be patterned through the DMD. This greatly simplified our targeted illumination module that the DMD surface can be directly imaged onto the sample through a tube lens and an objective, as opposed to alternative systems that requires dual-color excitation^5^ or more complicated holographic targeting^6^. For wide FOV SomArchon voltage imaging, large area excitation with a power density on the order of a few W/mm^2^ is necessary. While typically LEDs have insufficient power density, it can be easily satisfied with a multimode laser diode array because of the widefield nature of our DMD-based targeting strategy. Such light source would also avoid the undesirable speckle artifacts associated with targeting techniques that require coherent sources^6^. Additionally, in our system, the initial GFP channel structural imaging was performed with the same widefield microscope using an extra blue LED excitation. Even though this can also be performed using an additional two-photon microscope^5,6^, the integrated widefield microscope design described significantly reduces the cost and the complexity of the optical system and the software control. Since both GFP and SomArchon fluorescence were capture by the same camera, both images were automatically co-registered, further alleviating the issue of image registration and long-term system stability across multiple microscope modules. Overall, such simplified designs provide a cost-effective, easy to implement solution for large scale voltage imaging analysis of neural networks in behaving animals.

Single cell level imaging in living animals is always subject to fine movement due to metabolic, physiologic, and vascular changes, and a solution to such fine motion interference is through offline correction via image registration^24,25^. Targeting illumination only to cell membranes using holographic projections however is sensitive to translational movement due to the restricted area of illumination, which introduces additional challenge on the preparation of animal subjects, and the design of the behavioral tasks. The DMD-based widefield targeting strategy described here alleviate some of these concerns, where the illumination window size can be easily adjusted to accommodate fine biological motion. For example, increasing the region of targeted illumination to capture the morphological details of a neuron allows for fine translational movement that can be effectively corrected offline after image acquisition. Single photon widefield imaging is also more amenable to axial motions due to the lack of optical sectioning^18^, which allows for continuous recording of signal during subtle fluctuations in axial positions. While such advantage is retained to some degree over the illuminated regions with DMD-based targeted illumination, it is less so when using holographic projections^6^. Techniques with confined excitation volume limited to narrow z-axis profiles, such as two-photon microscopy^26^, are also more sensitive to image motion expected from behaving animals^27^.

To estimate SomArchon fluorescence quality, we calculated spike SBR. The baseline used in this SBR calculation contains both biological subthreshold membrane voltage fluctuations and SomArchon intrinsic fluorescent shot noise. Neurons in intact neural circuits, especially in the awake brain, receive heterogenous synaptic inputs and exhibit distinct membrane biophysical properties, which lead to variation in subthreshold membrane voltage fluctuations that are difficult to estimate. Thus, by itself, spike SBR estimation for each neuron cannot fully capture the quality of SomArchon signal contrasts, and represents an underestimation of SomArchon performance as illustrated with our spiking neuron models. However, spike SBRs of the same neuron when compared under the two illumination conditions can provide a quantitative measure of the fluorescence signal quality, providing direct experimental evidence that targeted illumination significantly improve the quality of SomArchon voltage imaging. Since spike SBR is a key consideration for spike detection, the fact that we detected more spikes under targeted illumination condition further demonstrates the improvement of SomArchon voltage signal quality.

## Acknowledgements

We thank members of the Han Lab for technical support.

## Author Contributions

S.X., E.L., H.J.G, and J.S. performed all experiments. S.X. and E.L. produced the simulated data model. E.L., P.F., and Y.W. analyzed the data. R.M. provided surgical expertise and H.T. consulted on imaging data analysis. H.M. provided *in vitro* resources and J.M. consulted on imaging system design. X.H. supervised the study. S.X, E.L., H.J.G, and X.H. wrote the manuscript. All authors edited the manuscript.

## Declaration of Interests

The authors declare no competing interests.

## Financial Disclosure

X.H. acknowledges funding from NIH (1R01MH122971, 1R01NS115797, R01NS109794, 1R34NS111742), NSF (CBET-1848029, DIOS-2002971). J.M. and X.H. acknowledges funding from NIH (R01EB029171), and E.L. acknowledges funding from Boston University Center for Systems Neuroscience. J.S. is supported by a training grant from the NIH/NIGMS (5T32GM008541-23). The funders had no role in study design, data collection and analysis, decision to publish, or preparation of the manuscript.

## Methods

### Simulated data theory for widefield fluorescence imaging

We consider the problem of widefield fluorescent imaging in a mouse brain in the context of imaging through scattering media within the forward scattering limit. In our model (Fig. S1), an incoherent source located at plane *z* = 0 is embedded at depth *z* = *z*_*t*_inside a scattering medium, whose scattering properties are characterized by the scattering phase function *p*(**ŝ**), mean scattering length *l*_*s*_, and anisotropic factor *g* ≈ 1. The image of the scattered light field from the source is relayed by a unit magnification 4*f*system and recorded by a detector that’s conjugate to the plane *z* = *z*_*s*_ in the sample space. Within the scattering medium, light propagation can be characterized in terms of radiance ℛ (*z*, **ρ**,**ŝ**) using a simplified radiative transport equation by invoking small angle approximation^22^:

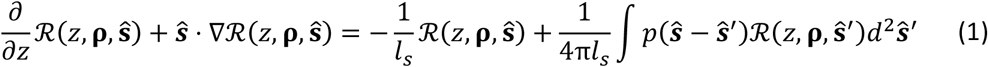

where(**ρ**, *z*) = (*x, y, z*) is the 3D position vector, **ŝ** = (*θ*_*x*_, *θ*_*y*_, 0) is a unit direction vector parameterized by the two angles assumed to be small, where *θ*_*x,y*_≈ 0.

To solve Eq. (1), it is necessary to establish a boundary condition, which, in our case, can be expressed as an isotropic emitter with intensity distribution *I*_0_(**ρ**_0_) located at axial position *z*_0_ = 0

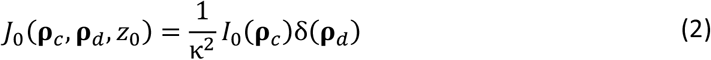

where *κ* =*n/ λ, n* is the refractive index, *λ* is the wavelength. Note that here we expressed the boundary condition in terms of mutual coherence function *J*(**ρ** _*c*_, **ρ** _*d*_, *z*) =⟨*E*(**ρ** _*+*_, *z*) *E*^*^ (**ρ** _−_, *z*) ⟩, where **ρ** _±_ = **ρ** _*c*_ ±**ρ** _*d*_ /2, instead of radiance ℛ (*z*, **ρ**,**ŝ**). This is because for an imaging system, we are more interested in the propagation of mutual coherence since it characterizes correlations of light fields between pairs of points that contribute to the final fluorescence intensity. This quantity, under paraxial limit, is related to the radiance of light field as

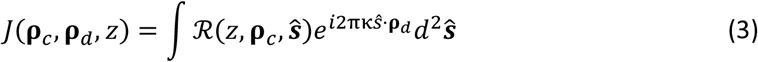

Eq. (1) together with the boundary condition Eq. (2) can be solved analytically using double Fourier transform^22^. We can therefore find the mutual coherence function at the surface of the scattering medium *z* = *z*_*t*_as

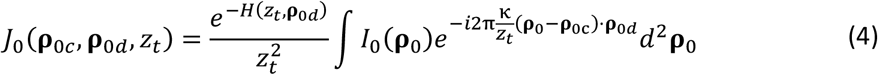

where 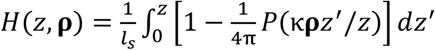, and *P****(q)*** = ∫*p****(s)****e*^*i*2π***s.q***^*d**s*** is the Fourier transform of the scattering phase function. This light field can be further propagated through a 4*f*imaging system, resulting in the measured intensity at the detector plane as

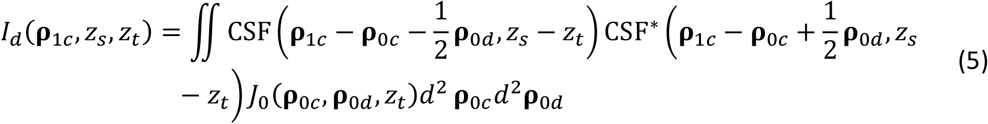

where 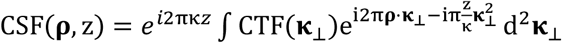 is the 3D coherent spread function, 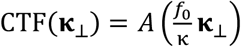 is the in-focus coherent transfer function, *A*(**ξ**) is the microscope aperture, and *f*_*0*_ is the focal length of the imaging lenses^16^. Note that here we assumed unit magnification and refractive index of the medium *n=*1 From Eq. (4) and Eq. (5), using the definition of optical transfer function we therefore have the 3D scattering optical transfer function (SOTF) for imaging a fluorescent object embedded in scattering media as:

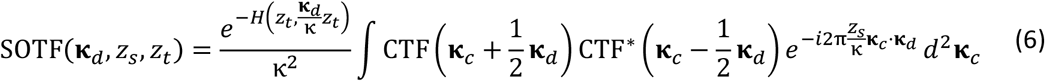

Eq. (6) is the main results that we use for simulating widefield neuronal imaging, the interpretation of which is that the propagation of mutual coherence can be simply considered as free space propagation with an additional attenuation factor 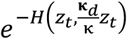 due to scattering. Note that this result not only holds for imaging of fluorescent signals in the detection path, but can also be applied to delivering illumination patterns onto a scattering sample in the excitation path (i.e., targeted illumination).

Biological tissues such as the brain are typically characterized by strong forward scattering where *g* ≈ 1, where the distribution of scattering angles follows the Henyey-Greenstein phase function^28^:

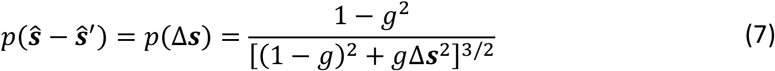

Assuming a circular microscope aperture of radius *r*, substituting Eq. (7) into Eq. (6) and using the Stokseth approximation of free space 3D optical transfer function (OTF)^29^, we arrive at the analytical solution of the 3D SOTF:

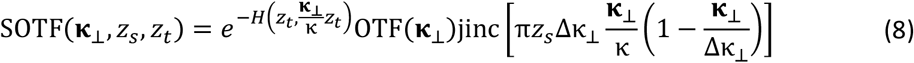

where Δ κ _⊥_ = 2*NA/λ,NA* = *r/f*_0_ is the numerical aperture of the system, and

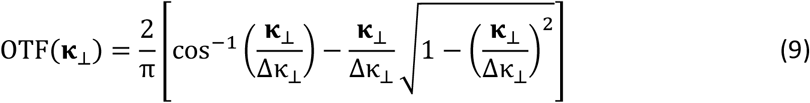

is the in-focus free space OTF. With Eq. (8), we can calculate the detected image or projected pattern simply by filtering the original object/pattern in frequency space using the corresponding SOTF.

### Simulation of widefield illumination versus targeted illumination conditions

Using the theoretical model developed above, we compared the background fluorescence signals generated using widefield and targeted illumination. We estimated the reduction of background fluorescent signals from non-targeted SomArchon expressing neurons, or in other words, signal cross-contamination, with the use of targeted illumination compared to standard widefield illumination. For simplicity, here we only modeled a pair of neurons that are separated by a distance *D*_*x*_ laterally and *D*_*z*_ axially (Fig. S2 and Fig. S3). Each neuron was assumed to be a 15 µm diameter uniformly fluorescent sphere. For widefield illumination, the entire FOV was illuminated equally. For targeted illumination, only a 15 µm circular ROI was projected onto the sample centered at the location of the neuron of interest. Although both neurons were imaged onto the camera, only the targeted one contained the signal, and the contribution from the other non-targeted neuron within the ROI of the targeted neuron (the red circle in Fig. S2 d-i and Fig. S3 c-h) was considered as background (or crosstalk).

In the simulation, we assumed the imaging system has unit magnification and *NA* = 0 4. The excitation and emission wavelength are *λ* _*ex*_ = 637nm, *λ* _*em*_ = 670nm respectively, with corresponding tissue anisotropic factor *g* _637nm_ = 0 89, *g* _670nm_ = 0.90, and mean scattering length *l*_*s*,637nm_ =110 μm, *l*_*s*,670nm_ =119 μm ^30^. Two different scenarios for optical voltage imaging were considered, namely *in vitro* imaging in 2D neuronal cell culture and *in vivo* imaging in a mouse brain.

For *in vitro* imaging in cultured neurons, since it typically consists of a monolayer of cells, we therefore assumed the two neurons are at the same depth *z* _*t*_= 0 (Fig. S2 a,b) with no tissue scattering. By varying lateral distance *D*_*x*_, we plotted the amount of crosstalk induced by the non-targeted neurons in Fig. S2 c. Both widefield illumination and targeted illumination introduce similar amount of crosstalk, as confirmed by our *in vitro* imaging experiments. Note that in reality, these two neurons should not overlap in space and should have a separation at least *D*_*x*_ =15 µm (although *D*_*x*_ < 15 µm is still plotted for completeness), in which case the amount of crosstalk is close to 0. Therefore, we expect very little benefit of using targeted illumination for reducing crosstalk in *in vitro* imaging. However, targeted illumination still pertains certain advantages over widefield illumination in terms of photobleaching and SBR because of the reduction of stray light and non-specific background signals.

For *in vivo* imaging in a mouse brain, we assumed that the targeted neuron was located at depth *z*_*t*_ =100 µm (see Fig. S3 a,b) inside the tissue with scattering properties given above. The amount of crosstalk at positions with varying *D*_*x*_ and *D*_*z*_ are plotted in Fig. 1c. In this case, targeted illumination results in much higher reductions in crosstalk, with most significant effects when out-of-focus (*D*_*z*_ ≠ 0). Example images of the non-targeted neuron at a defocus distance *D*_*z*_ = 15 µm with varying lateral displacement *Dx* = 0 µm, 10 µm, 20 µm are given in Fig. S3 c-h, where one can see much lower intensity from the non-targeted neuron, with the crosstalk under targeted illumination only at 76%, 44% and 3.8% of the values under widefield illumination.

Note that here our simulation only considers a pair of neurons, so the induced crosstalk values are relatively low. For *in vivo* imaging where a much higher number of neurons are labeled, the signal cross-contaminations can be introduced by tens or hundreds of neurons collectively. In this case, the background with widefield illumination would be more detrimental as to render the in-focus neuron visually indiscernible (see Fig. 1d-g), which could further necessitate the use of targeted illumination.

### Membrane voltage simulation

The membrane voltage *v* was of Izhikevich-type and defined as follows:

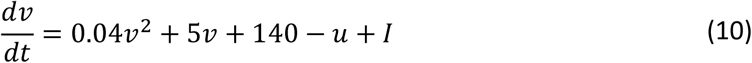

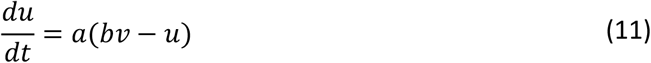

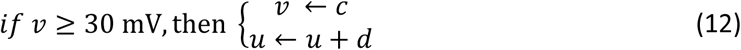

The two coupled differential equations were numerically solved using the Euler method with 1 ms step size, modeling a 1KHz sampling rate. For the parameters, we chose: a = 0.02, b = 0.2, c = -55 mV, d = 2 as often used^31^. The input to the neuron was composed of a fixed input current to each neuron (2.8 mv) and gaussian current noise (standard deviation = 2 mV).

To estimate the spike SBR (see details below), we divided the spike amplitude (here defined as -50 mV to 30 mV = 80 mV) by the baseline voltage fluctuations, including both the biological (dynamic) noise variance and the added measurement gaussian white noise variance. SNR was defined as the spike amplitude divided by the added gaussian white noise variance only.

### Cell cultures

Rat cortical neuron cultures were dissociated from E18 rat embryos (Charles River) and plated on coverslips coated with poly-D lysine (Millipore Sigma cat # P2636) at 0.1 mg/mL in 0.1M borate buffer (pH 8.5), and bathed with plating medium containing DMEM/F12 (Gibco cat. # 21331020) supplemented with 10% Heat Inactivated FBS (R&D systems cat. # S11150), 5% Heat Inactivated Horse Serum (Thermo Fisher Scientific cat. # 26050070), 1% Penicillin/Streptomycin (Thermo Fisher Scientific cat. # 15140122), 397 µM L-Cysteine hydrochloride (Millipore Sigma cat. # C1276), and 2 mM L-Glutamine (Thermo Fisher Scientific cat. # 35050061) (O’Connor 2020). 24 hours after plating, cells were switched to a feeding medium containing NBM (Gibco cat. # 21103049) supplemented with 1% Heat Inactivated Horse Serum (Thermo Fisher Scientific cat. # 26050070), 2% NeuroCult SM1 supplement (Stimcell Technologies cat. # 05711), and 1.4% penicillin/streptomycin (Thermo Fisher Scientific cat. # 15140122) and 800 µM L-Glutamine. 11 days later, 5-fluoro-2-deoxyuridine (Millipore cat. # 343333) was added at a concentration of 4 µM to prevent glial cell overgrowth. 50% of the cell culture medium was exchanged every 3 days. Neurons were transduced with 0.25 µL of AAV9-syn-SomArchon per well in 0.25 mL of feeding media, 3-4 days after plating. Cells were imaged 14-16 days after plating, in an imaging buffer containing 145 mM NaCl, 2.5 mM KCl, 10 mM glucose, 10 mM HEPES, 2 mM CaCl_2_, and 1 mM MgCl_2_, pH 7.4.

### Animal surgical procedures

All procedures involving animals were approved by the Boston University Institutional Animal Care and Use Committee (IACUC). C57BL/6 adult female mice (3-6 months old on the day of recording) were used in this study. Mice were surgically implanted with an imaging chamber and a head-plate as described previously^3^. AAV-syn-SomArchon was injected either through an infusion cannula attached to the window after the surgery, or injected during the surgery.

### Custom widefield optical imaging setup

We customized a dual color epi-fluorescence fluorescence microscope, which used a 470 nm LED (Thorlabs, M470L3) for GFP fluorescence excitation, and a 637 nm fiber-coupled laser (Ushio America Inc., Necsel Red-HP-FC-63x) for SomArchon fluorescence excitation. The two illumination channels were combined using a dichromatic mirror (Thorlabs, DMLP550R) and subsequently directed onto the sample. The generated fluorescent signal was epi-collected by a microscope objective (Nikon, 40×/0.8NA CFI APO NIR) and imaged onto a camera (Hamamatsu, ORCA-Lightning C14120-20P) with a 175 mm tube lens. A combination of excitation filter, dichromatic mirror, and emission filter (Semrock, LF405/488/532/635-A-000) was used to separate fluorescent signals from the excitation light.

To pattern the illumination in the SomArchon imaging channel, the output of the 637 nm multimode laser was collimated (Thorlabs, F950SMA-A), expanded (Thorlabs, BE02M-A), and directed onto a DMD (Vialux, V-7000 VIS) at approximately 24° to its surface normal. The DMD was further imaged onto the sample with a 175 mm lens and the objective, so that only sample regions corresponding to the ‘on’ pixels of DMD were illuminated. The axial position of the DMD was adjusted so that it is conjugate to the camera, and an additional affine transform was estimated to register the pixels between the DMD and the camera. The DMD was controlled using custom Matlab script based on Vialux ALP-4.2 API.

During each imaging session, a GFP fluorescence image was first taken for illumination target identification, where a small rectangular ROI was manually selected for each individual neuron to be imaged. A binary illumination mask was then generated based on all the selected ROIs and uploaded to the DMD for illumination targeting. SomArchon voltage imaging was performed at 500 Hz, with 2 × 2 pixel binning, resulting in an imaging area of 1152 × 576 pixels on the sCMOS camera sensor, corresponding to a 360 × 180 µm^2^ FOV at the sample. To estimate sCMOS camera dark level and intrinsic noise, videos were collected with the camera set to the same acquisition parameters as during regular imaging experiments, but without light exposure (500 Hz, 2 × 2 pixel binning, 1152 × 576 pixels imaging area). The sensor dark level was estimated to be 767.7, with an intrinsic noise of 12.6 (standard deviation) per pixel.

### Data analysis

All imaging data were acquired by HCImage software (Hamamatsu), and further processed using MATLAB (Mathworks) offline.

### Neuron ROI selection

All data analysis was performed offline in Matlab 2019b or 2020a. SomArchon fluorescence images were first motion corrected using a pairwise rigid motion correction algorithm as described previously^24^. For targeted illumination recordings, each ROI was centered on a neuron of interest, with the ROI size slightly greater than the outline of the neurons, so that motion correction can be performed on each targeted ROI that had distinguishable features identifiable by the algorithm. After motion correction, we manually selected ROIs corresponding to individual neurons, based on the average SomArchon fluorescence image during the first recorded trial. ROIs were cross-referenced by comparing SomArchon fluorescence with the stable EGFP fluorescence. The identified neurons were then applied to all subsequent trials in the same recording session. SomArchon fluorescence traces were then extracted for each neuron by averaging all the pixels within the neuron across the entire experiments. For direct comparison of SomArchon fluorescence of the same neurons between widefield and target illumination conditions, the same neuron ROIs were used for both recording conditions. Trace time segments with sharp, drastic changes in fluorescence (e.g. due to motion) were detected as outliers and excluded from further analysis in both the widefield and targeted illumination analysis. Specifically, for the outlier detection we applied the generalized extreme Studentized deviate test on the moving standard deviation values using a sliding window of ±60 ms on spike-removed traces (see Method Section spike detection and spike SBR calculation). In some cases, not all time points during the period of an artefact were marked as outliers. Time points between outliers (< 3 data points) were therefore interpolated. To remove further artefacts, we excluded time points that were 6 standard deviations outside the trace fluorescence distribution. Time points between and around the detected outliers were also removed (±350 ms) as those periods often coincided with extended animal motion artefacts.

### Fluorescence decay estimation

To estimate SomArchon fluorescence decay, we first removed spikes by applying a median filter (window of 51 frames), and then subtracted the camera dark level (measured as 767.7). We calculated fluorescence decay as the ratio of the mean fluorescence intensity during the first 600 ms and that during the last 600 ms for each trial, and then averaged across all trials. In cultured neurons, we detected a drastic fluorescence drop within the first couple seconds of recording, likely mainly due to bleaching of autofluorescence unrelated to SomArchon, thus we excluded the first trial from subsequent analysis for culture neuron analysis.

### Spike detection and spike SBR calculation

To separate spikes from subthreshold voltage fluctuations, we first generated a “Smoothed Trace” (ST) by averaging the fluorescence trace using a moving window of ±100 frames. To estimate baseline fluctuation, we first removed potential spikes by replacing any fluorescence values above ST with the corresponding values of ST. The amplitude of the baseline fluctuation was then estimated as 2 times the standard deviation of the resulting trace, since half of the fluctuations were removed in the spike removal step described above. For spike SBR estimation, we also subtracted the camera intrinsic noise (standard deviation = 12.6 per pixel) from the trace noise to obtain camera-independent estimates.

For spike detection, we first removed small subthreshold rapid signal changes by replacing the fluorescence below ST with corresponding values of ST. The derivative of the resulting trace was then used for spike detection, where spikes were identified as the time points above 4.5 times of the standard deviation of the resulting derivative trace. Spike amplitude was calculated as the peak fluorescence for each spike minus the mean of the fluorescence during the three time points before spike onset. Spike SBR was calculated as spike amplitude divided by the amplitude of baseline fluctuations described above.

### Pearson correlation analysis

Pearson cross-correlation was performed using the Matlab functions *corrcoef* and *xcorr*, for both Vm-Vm and spike-spike correlation analysis. To calculate spike-spike correlation, spike vectors were smoothed over a ±10 ms time window for each spike before applying correlation analysis. To calculate Vm-Vm correlation, we removed spikes by replacing 3 data point centered at each identified spike times with the adjacent values that largely eliminated the contribution of spikes in Vm-Vm correlation analysis.

### Statistical analysis

Paired student’s t-tests were used for comparisons involving the same neurons between the targeted illumination condition and the widefield illumination condition. A Kolmogorov-Smirnov test was used to test the difference of cross-correlation over distance between targeted illumination and widefield illumination conditions. For Kolmogorov-Smirnov test, the data in each of the two conditions were first sorted by distance before comparing. A Friedman’s test, 2 factor non-parametric ANOVA, was used to compare the difference between the average correlations of each bin in Fig. 3.

### Data and software availability statement

Codes used for data analysis is available on our lab website and Github repository: https://www.bu.edu/hanlab/resources/ and https://github.com/HanLabBU

## Notes

### Competing Interest Statement

The authors have declared no competing interest.

## References

1. Gong, Y. et al. High-speed recording of neural spikes in awake mice and flies with a fluorescent voltage sensor. Science 350, (2015).

2. Abdelfattah, A. S. et al. Bright and photostable chemigenetic indicators for extended in vivo voltage imaging. Science 365, (2019).

3. Piatkevich, K. D. et al. Population imaging of neural activity in awake behaving mice. Nature 574, (2019).

4. Villette, V. et al. Ultrafast two-photon imaging of a high-gain voltage indicator in awake behaving mice. Cell 179, (2019).

5. Adam, Y. et al. Voltage imaging and optogenetics reveal behaviour-dependent changes in hippocampal dynamics. Nature 569, (2019).

6. Fan, L. Z. et al. All-optical electrophysiology reveals the role of lateral inhibition in sensory processing in cortical layer 1. Cell 180, (2020).

7. Bando, Y., Grimm, C., Cornejo, V. H. & Yuste, R. Genetic voltage indicators. BMC Biology 17, (2019).

8. Xu, Y., Zou, P. & Cohen, A. E. Voltage imaging with genetically encoded indicators. Current Opinion in Chemical Biology 39, (2017).

9. Kannan, M., Vasan, G. & Pieribone, V. A. Optimizing strategies for developing genetically encoded voltage indicators. Frontiers in Cellular Neuroscience vol. 13 (2019).

10. Beck, C., Zhang, D. & Gong, Y. Enhanced genetically encoded voltage indicators advance their applications in neuroscience. Current Opinion in Biomedical Engineering 12, (2019).

11. Knöpfel, T. & Song, C. Optical voltage imaging in neurons: moving from technology development to practical tool. Nature Reviews Neuroscience 20, (2019).

12. Peng, L., Xu, Y. & Zou, P. Genetically-encoded voltage indicators. Chinese Chemical Letters 28, 1925–1928 (2017).

13. Ma, Y., Bayguinov, P. O. & Jackson, M. B. Optical studies of action potential dynamics with hVOS probes. Current Opinion in Biomedical Engineering 12, (2019).

14. Piatkevich, K. D. et al. A robotic multidimensional directed evolution approach applied to fluorescent voltage reporters. Nature Chemical Biology 14, (2018).

15. Abdelfattah, A. S. et al. A general approach to engineer positive-going eFRET voltage indicators. Nature Communications 11, (2020).

16. Mertz, J. Introduction to optical microscopy. Introduction to Optical Microscopy (2019). doi:10.1017/9781108552660.

17. Wu, J. et al. Kilohertz two-photon fluorescence microscopy imaging of neural activity in vivo. Nature Methods 17, (2020).

18. Mertz, J. Optical sectioning microscopy with planar or structured illumination. Nature Methods 8, (2011).

19. Harris, K. D., Quiroga, R. Q., Freeman, J. & Smith, S. L. Improving data quality in neuronal population recordings. Nature Neuroscience 19, (2016).

20. Lim, S. T., Antonucci, D. E., Scannevin, R. H. & Trimmer, J. S. A novel targeting signal for proximal clustering of the Kv2.1 K+ channel in hippocampal neurons. Neuron 25, (2000).

21. Xiao, S., Tseng, H. A., Gritton, H., Han, X. & Mertz, J. Video-rate volumetric neuronal imaging using 3D targeted illumination. Scientific Reports 8, (2018).

22. Ishimaru, A. Wave propagation and scattering in random media. Wave Propagation and Scattering in Random Media (1999). doi:10.1109/9780470547045.

23. Brivanlou, I. H., Dantzker, J. L. M., Stevens, C. F. & Callaway, E. M. Topographic specificity of functional connections from hippocampal CA3 to CA1. Proceedings of the National Academy of Sciences of the United States of America 101, (2004).

24. Pnevmatikakis, E. A. & Giovannucci, A. NoRMCorre: An online algorithm for piecewise rigid motion correction of calcium imaging data. Journal of Neuroscience Methods 291, (2017).

25. Mohammed, A. I. et al. An integrative approach for analyzing hundreds of neurons in task performing mice using wide-field calcium imaging. Scientific Reports 6, (2016).

26. Helmchen, F. & Denk, W. Deep tissue two-photon microscopy. Nature Methods 2, (2005).

27. Griffiths, V. A. et al. Real-time 3D movement correction for two-photon imaging in behaving animals. Nature Methods 17, (2020).

28. Jacques, S. L. Optical properties of biological tissues: A review. Physics in Medicine and Biology 58, (2013).

29. Stokseth, P. A. Properties of a defocused optical system. Journal of the Optical Society of America 59, (1969).

30. Yaroslavsky, A. N. et al. Optical properties of selected native and coagulated human brain tissues in vitro in the visible and near infrared spectral range. Physics in Medicine and Biology 47, (2002).

31. Izhikevich, E. M. Simple model of spiking neurons. IEEE Transactions on Neural Networks 14, (2003).

