## Supplementary material for "Large-scale voltage imaging in the brain using targeted illumination": Supplmenental materials

### Supplementary Materials:

### Supplementary Figures:

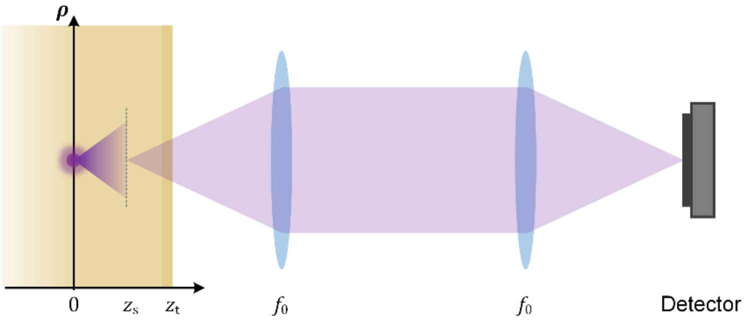


***Figure S1:*** *Theoretical model for simulating fluorescent imaging in a mouse brain: an incoherent source located at z = 0 is at distance $z_t$ away from the surface of the scattering media, $z_s$ away from the focal plane of the microscope. The fluorescent signal is imaged by a unit magnification $4f$ system to the detector.*

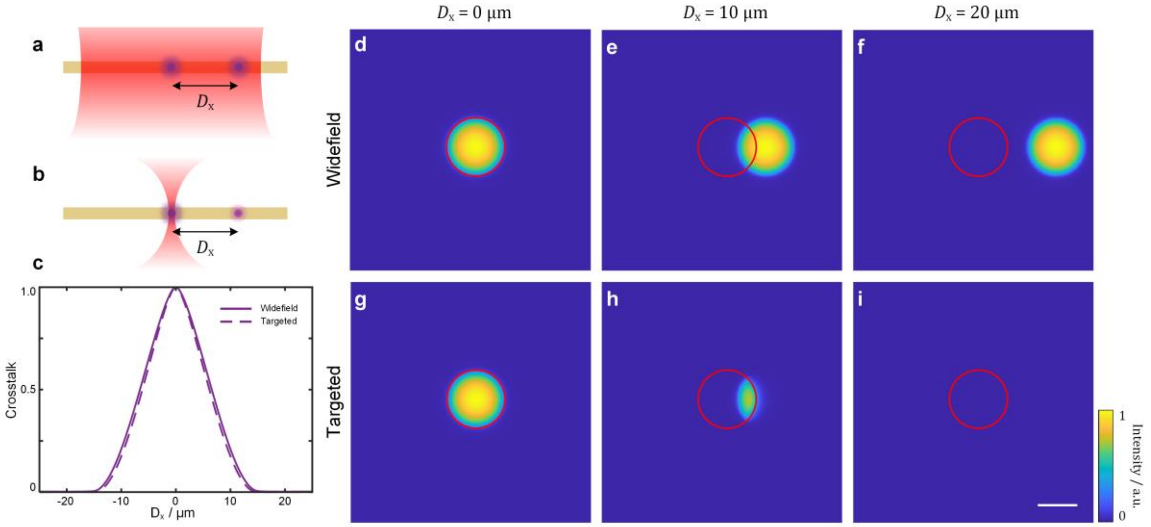


***Figure S2****: Simulation results of in vitro optical voltage imaging in neural cell cultures. (****a,b****) Illustration of in vitro imaging a pair of neurons using widefield (****a****) and targeted (****b****) illumination respectively. (****c****) Comparison of signal crosstalk induced by the non-targeted neuron at varying lateral distance $D_x$. (****d-i****) Simulated images of the non-targeted neuron at lateral distance $D_x = 0$ µm, 10 µm and 20 µm under widefield (****d-f****) and targeted (****g-i****) illumination. Red circles represent the ROI of the targeted neuron. Scale bar, 10 µm. a.u., arbitrary unit.*

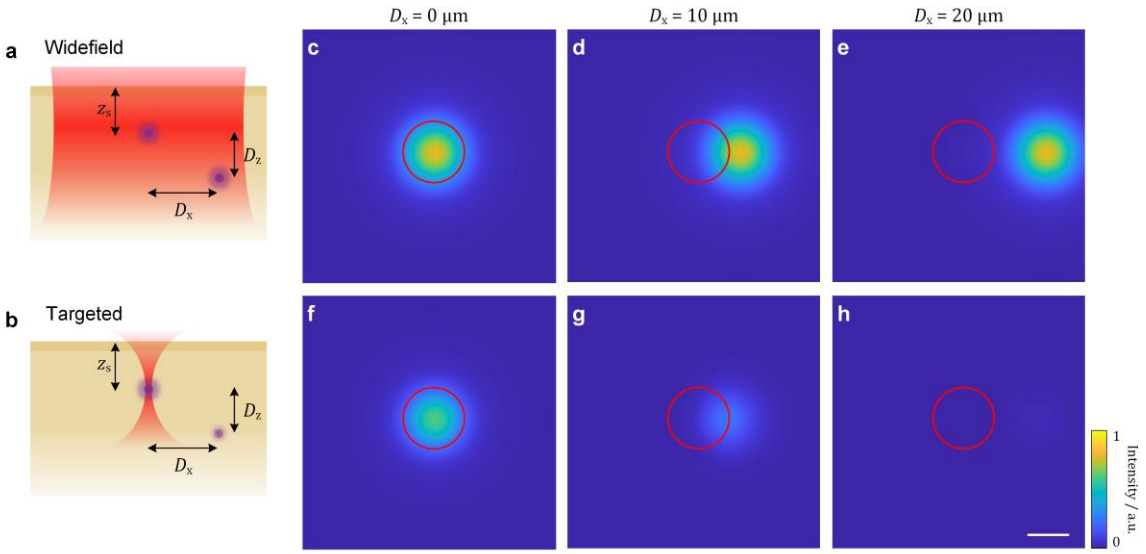


***Figure S3:** Simulation results of in vivo optical voltage imaging in a mouse brain. (**a,b**) Illustration of in vivo imaging a pair of neurons using widefield (**a**) and targeted (**b**) illumination respectively. (**c-h**) Simulated images of the non-targeted neuron at fixed axial distance $D_z = 15$ µm and varying lateral distance $D_x = 0$ µm, 10 µm and 20 µm under widefield (**c-e**) and targeted (**f-h**) illumination. Red circles represent the ROI of the targeted neuron. Scale bar, 10 µm. a.u., arbitrary unit.*

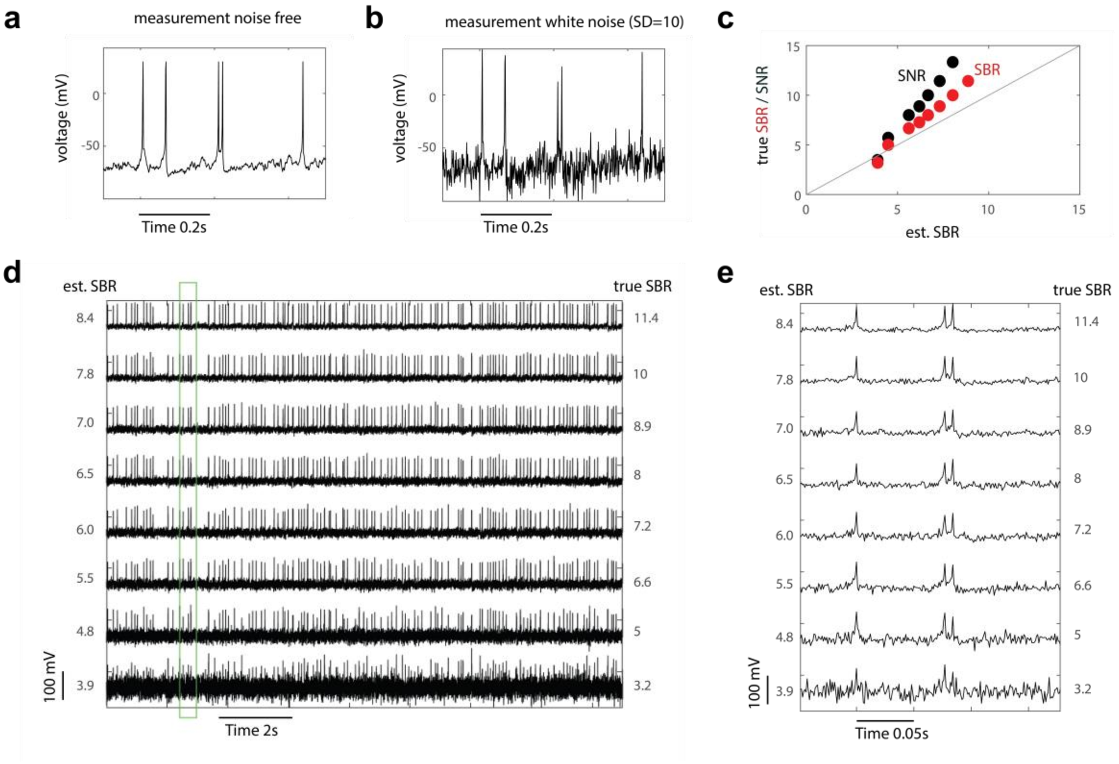


***Figure S4***. Stimulations of an integrate-and-fire neuron, with varying levels of measurement noise. (**a**) Simulated membrane voltage traces without any measurement noise added. (**b**) Same simulated membrane voltage trace with Gaussian white noise added (SD = 10). (**c**) Comparison of the estimated spike SBR using the same algorithm used for experimental data, and the theoretical ground truth SBR (red) and ground truth SNR (black). The ground truth SBR is defined as the spike amplitude (from base -50 mV to spike peak +30 mV = 80 mv) divided by variation of Vm that contains both the added noise and the biological noise. SNR calculation is without the biological noise. (**d**) The same simulated membrane voltage trace with different levels of added white noise and the corresponding estimated and ground truth SBR. (**e**) Zoomed-in version of (**d**) as indicated by the green box.

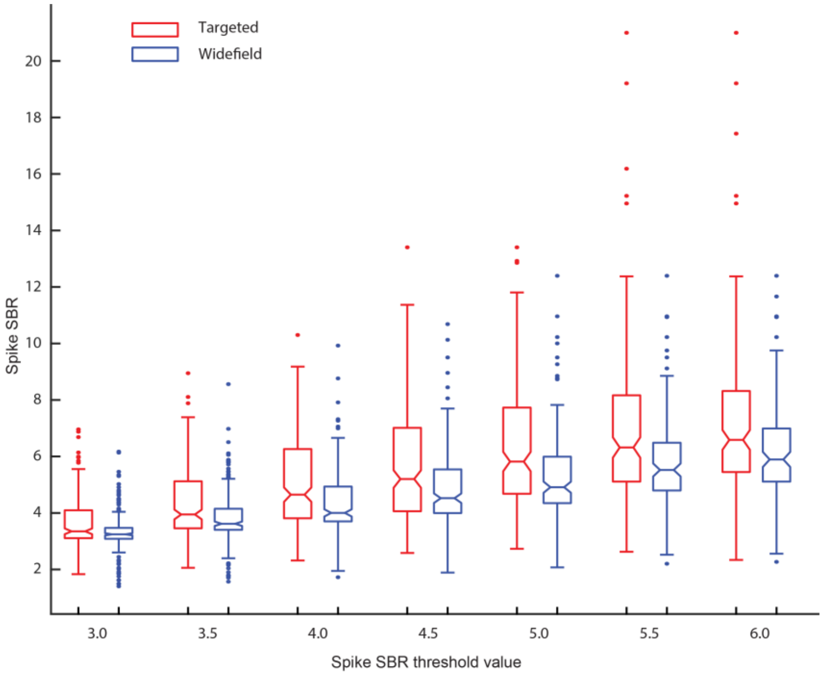


***Figure S5:*** *Targeted illumination improves spike SBR regardless of spike detection threshold used. Spike SBR for the spikes detected using different spike detection threshold values under targeted illumination (red) and widefield illumination (blue) conditions. Increasing spike detection threshold correspondingly increased spike SBR of the detected spikes, for both recording conditions. However, regardless of the spike detection threshold chosen, the spikes detected under targeted illumination conditions have greater spike SBR than that detected under widefield illumination conditions. Refer to Table S3 for statistical tests. Each data point in the plot represents median SBR for all neurons, and error bars represent the standard deviation. For all boxplots, the box indicates the median (middle line), 25th (Q1, bottom line), 75th (Q3, top line) percentiles, and the whiskers are Q1-1.5*(Q3-Q1), and Q3+1.5*(Q3-Q1). Outliers that exceed these values are shown as dots.*

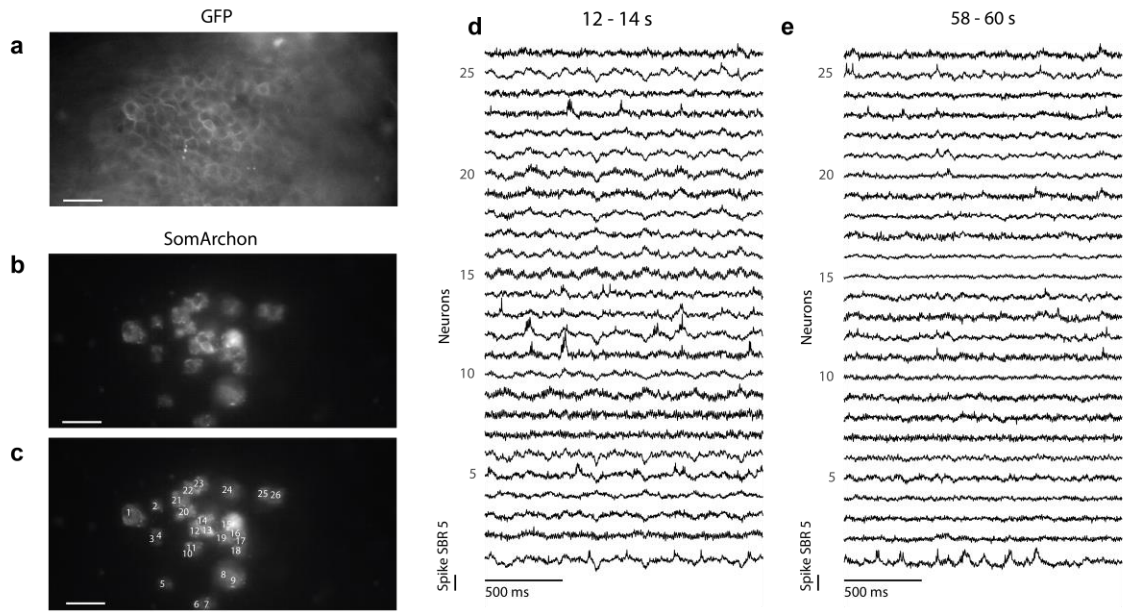


***Figure S6:*** *An example 80-second long recording session with 25 simultaneously recorded CA1 neurons using targeted illumination in a behaving mouse. (**a-c**) SomArchon expressing CA1 neurons in the FOV, visualized via GFP fluorescence (**a**), SomArchon fluorescence with targeted illumination (**b**), and with each neuron labelled (**c**). Scale bar, 50 µm. (**d,e**) Example fluorescence traces of simultaneous recording of 25 neurons. Shown are the time segment 12-14 s (**d**) and 58-60 s (**e**) of the overall recordings.*

### Supplementary Tables:

| Binned Distance (µm) | Median | | p-Value | Confidence Interval | Degrees of Freedom | T-statistics | Std Dev |
|---|---|---|---|---|---|---|---|
| | Targeted | Widefield | | | | | |
| 0-29 | 0.16287 | 0.17023 | 0.29026 | -0.0084656, 0.028232 | 389 | 1.059 | 0.18431 |
| 30-59 | 0.10654 | 0.13304 | 0.48894 | -0.014956, 0.007158 | 693 | -0.69237 | 0.14836 |
| 60-89 | 0.094583 | 0.1239 | 0.98747 | -0.009993, 0.009835 | 801 | -0.015704 | 0.14303 |
| 90-119 | 0.077449 | 0.10864 | 0.36773 | -0.015469, 0.0057354 | 665 | -0.90135 | 0.13935 |
| 120-149 | 0.07688 | 0.11532 | 1.28E-06 | -0.030586, -0.013065 | 583 | -4.8934 | 0.10779 |
| 150-179 | 0.068209 | 0.091214 | 9.70E-10 | -0.02776, -0.014527 | 353 | -6.2845 | 0.0633 |
| >=180 | 0.049822 | 0.086531 | 2.42E-06 | -0.021796, -0.009123 | 325 | -4.7996 | 0.058156 |

***Table S1:*** *Summary of paired t-test statistics comparing Vm-Vm correlations under targeted illumination versus widefield illumination condition.*

| Binned Distance (µm) | Median | | p-Value | Confidence Interval | Degrees of Freedom | T-statistics | Std Dev |
|---|---|---|---|---|---|---|---|
| | Targeted | Widefield | | | | | |
| 0-29 | 0.04211 | 0.018057 | 4.25E-07 | 0.017471, 0.03878 | 242 | 5.1997 | 0.084319 |
| 30-59 | 0.028633 | 0.0087085 | 1.58E-11 | 0.016339, 0.029241 | 391 | 6.9454 | 0.064966 |
| 60-89 | 0.024633 | 0.010663 | 4.67E-12 | 0.016927, 0.02988 | 465 | 7.101 | 0.071146 |
| 90-119 | 0.015225 | 0.012588 | 3.54E-05 | 0.0086944, 0.024064 | 334 | 4.1926 | 0.071505 |
| 120-149 | 0.01583 | 0.008564 | 3.07E-05 | 0.0099877, 0.027337 | 297 | 4.2339 | 0.076091 |
| 150-179 | 0.015378 | -0.0018637 | 3.12 E-04 | 0.0064034, 0.021229 | 175 | 3.6784 | 0.04983 |
| >=180 | 0.0054598 | 0.0022514 | 0.24885 | -0.0063823, 0.024428 | 140 | 1.158 | 0.092524 |

***Table S2:*** *Summary of paired t-test statistics comparing spike-spike correlations under targeted illumination versus widefield illumination condition.*

| SBR Threshold | Median | | P-Value | Confidence Interval | Degrees of Freedom | T-Statistics | Std Dev |
|---|---|---|---|---|---|---|---|
| | Targeted | Widefield | | | | | |
| 3.0 | 3.3506 | 3.2448 | 1.13E-14 | 0.2379, 0.3866 | 225 | 8.2750 | 0.5673 |
| 3.5 | 3.9473 | 3.6182 | 0 | 0.4173, 0.6473 | 225 | 9.1202 | 0.8774 |
| 4.0 | 4.6486 | 4.0029 | 0 | 0.5839, 0.9020 | 223 | 9.2066 | 1.2077 |
| 4.5 | 5.2018 | 4.5228 | 6.73E-14 | 0.6735, 1.1108 | 206 | 8.0448 | 1.5955 |
| 5.0 | 5.8190 | 4.9125 | 1.50E-12 | 0.8653, 1.4651 | 162 | 7.6716 | 1.9391 |
| 5.5 | 6.3176 | 5.5225 | 1.46E-09 | 0.8629, 1.6151 | 129 | 6.5176 | 2.1674 |
| 6.0 | 6.5870 | 5.8941 | 2.47E-09 | 0.6596, 1.2436 | 119 | 6.4529 | 1.6154 |

***Table S3:*** *Summary of paired t-test statistics comparing SBRs of detected spikes under targeted illumination versus widefield illumination condition. Only neurons that had detectable spikes in both conditions were considered for the paired t-test.*
